# The dynamic landscape of fission yeast meiosis alternative-splice isoforms

**DOI:** 10.1101/045922

**Authors:** Zheng Kuang, Jef D. Boeke, Stefan Canzar

## Abstract

Alternative splicing increases the diversity of transcriptomes and proteomes in metazoans. The extent to which alternative splicing is active and functional in unicellular organisms is less understood. Here we exploit a single-molecule long-read sequencing technique and develop an open-source software program called SpliceHunter, to characterize the transcriptome in the meiosis of fission yeast. We reveal 17017 alternative splicing events in 19741 novel isoforms at different stages of meiosis, including antisense and read-through transcripts. Intron retention is the major type of alternative splicing, followed by “alternate intron in exon”. 887 novel transcription units are detected; 60 of the predicted proteins show homology in other species and form theoretical stable structures. We compare the dynamics of novel isoforms based on the number of supporting full-length reads with those of annotated isoforms and explore the translational capacity and quality of novel isoforms. The evaluation of these factors indicates that the majority of novel isoforms are unlikely to be both condition-specific and translatable but the possibility of functional novel isoforms is not excluded. Moreover, the co-option of these unusual transcripts into newly born genes seems likely. Together, this study highlights the diversity and dynamics at the isoform level in the sexual development of fission yeast.

## Introduction

Splicing is a fundamental process which removes intragenic non-coding regions (introns) and forms mature mRNAs for proper translation (Lee and Rio 2015; Wang and Burge 2008). It provides an important checkpoint and a posttranscriptional layer of gene expression control (Bentley 2014; Braunschweig et al. 2013; McGlincy and Smith 2008; Le Hir et al. 2003). One of the regulatory mechanisms is alternative splicing, which leads to multiple transcripts from the same gene (Wang et al. 2008). Alternative splicing (AS) is achieved mainly by different combinations of exons and introns or AS sites, producing a vast expansion of transcriptome diversity and potentially, protein diversity (Nilsen and Graveley 2010; Keren et al. 2010). AS is considered as a potent regulator of gene expression in multicellular organisms given the rationale that isoforms generated by AS are differentially regulated in different tissues or conditions (Wang et al. 2008). However, AS in unicellular organisms is less extensively explored. Important questions are: What is the complexity of single cell transcriptomes? How is AS differentially regulated across different conditions? Is AS even functionally relevant to eukaryotic microorganisms? Understanding these questions in unicellular organisms will extend our knowledge of AS, and in particular the origin of AS.

*S. cerevisiae* and *Sch. pombe* are the two mostly documented unicellular model organisms. Unlike *S. cerevisiae, Sch. pombe* exhibits many features observed in metazoans, including heterochromatin structure, RNA interference and importantly, prevalence of introns (Rhind et al. 2011). *S. cerevisiae* has ~280 intron-containing genes and on average, each gene has only one intron. On the contrary, >2200 genes in *Sch. pombe* are currently known to contain introns and half of these genes have multiple introns. Therefore, *Sch. pombe* is a more suitable unicellular model to study AS and transcriptome diversity than *S.cerevisiae*. Thousands of AS events have been identified recently under various conditions in WT and mutant cells via different techniques such as short read RNA-seq and lariat sequencing (Bitton et al. 2015; Awan et al. 2013; Stepankiw et al. 2015). RNA metabolism kinetics, including synthesis, splicing and decay rates have also been determined in vegetative fission yeast (Eser et al. 2015). A broad spectrum of AS types is observed, similar to multicellular organisms. These studies suggest prevalent AS in this unicellular organism. However, these findings are limited in defining isoforms, which represent complete structure and sequence of transcripts. Characterizing isoforms is critical to predicting the effects on the proteome. Whether these AS events are functionally relevant or condition-specific also remains poorly explored.

To address these questions, we exploited a single-molecular real-time (SMRT) sequencing technique based on the Pacific Biosciences (PacBio) platform (Eid et al. 2009) and developed SpliceHunter, a novel open-source software program to systematically explore the transcriptome in *Sch. pombe*. PacBio sequencing features very long reads, which are suitable for full-length cDNA sequencing. Recently, PacBio sequencing has been used to examine the transcriptome in multiple metazoan organisms, such as chicken and human, and multiple plants, and identified thousands of annotated and novel isoforms (Sharon et al. 2013; Martin et al. 2014; Au et al. 2013; Tilgner et al. 2014; Minoche et al. 2015; Thomas et al. 2014; Dong et al. 2015). To systematically characterize isoforms and explore their differential regulations in fission yeast, we performed time-course isoform-level profiling during the meiosis of *Sch. pombe*. Meiosis is a well-documented process with a transition from vegetative growth to sexual differentiation. The transcriptome is dramatically reshaped and multiple AS events have been observed (Mata et al. 2002; Wilhelm et al. 2008). Here, we report 28379 isoforms occurring at different stages of meiosis, along with 17017 novel AS events in 19741 isoforms (Table S1). We estimated the abundance of individual isoforms using the numbers of supporting reads as proxy and analyzed their dynamics during meiosis. Although many isoforms are unlikely to produce meaningful proteins because of frameshifts and early termination, some novel isoforms and corresponding annotated isoforms show differential temporal patterns during meiosis and can generate distinct proteins. Additionally, we examined the conservation of the hypothetical proteins and evaluated their potential to form stable structures through a computational prediction of secondary and tertiary protein structure (Källberg et al. 2012). Together, this study for the first time summarizes the diversity and dynamics of the transcriptome at the isoform level during *Sch. pombe* meiosis and presents examples of AS isoforms that are condition-specific and potentially functionally relevant.

## Results

### Experimental design, workflow and definition of AS types

To explore the diversity and dynamics of alternative splicing in *Sch. pombe*, we collected 6 time points of WT cells during meiosis from 0 to 10 hours at 2 hour intervals (Figure S1). PolyA RNA was purified, reverse-transcribed to cDNA and sequenced with a PacBio sequencer. 5 SMRT cells were used for two RNA replicates of each time point. Raw reads were clustered and polished using the Iso-Seq pipeline and high-quality (HQ), full-length (FL), polished consensus sequences (referred to as Isoseq reads in the following) were output (Rhoads and Au 2015). We developed SpliceHunter, a novel software program that detects, quantifies, and compares complex splicing patterns along novel isoforms inferred from Isoseq reads. SpliceHunter reports the number of supporting reads of isoforms per sample or time point and predicts protein sequences. Based on the Iso-Seq algorithm, each Isoseq read is associated with multiple FL CCS (circular-consensus) reads and non-FL CCS reads. FL CCS reads are defined by co-existence of 5’ and 3’ adaptors and polyA tail. Therefore, we use the number of FL CCS reads as proxy for abundance and dynamics analysis.

After mapping reads to a reference genome, SpliceHunter assigns reads to annotated or novel transcription units (TUs) (Figure 1A, step 1) according to well-defined and adjustable criteria. After collapsing compatible reads to isoforms, each isoform is compared to the previously annotated exon-intron structure to detect alternative splicing events (Figure 1A, step 2, 3), antisense, and read-through transcripts. Finally, the effect of alternative splicing events on the encoded protein sequence is analyzed (Figure 1A, step 4), including their combined shift in the reading frame. See Methods for a detailed description of SpliceHunter.

**Figure 1.**
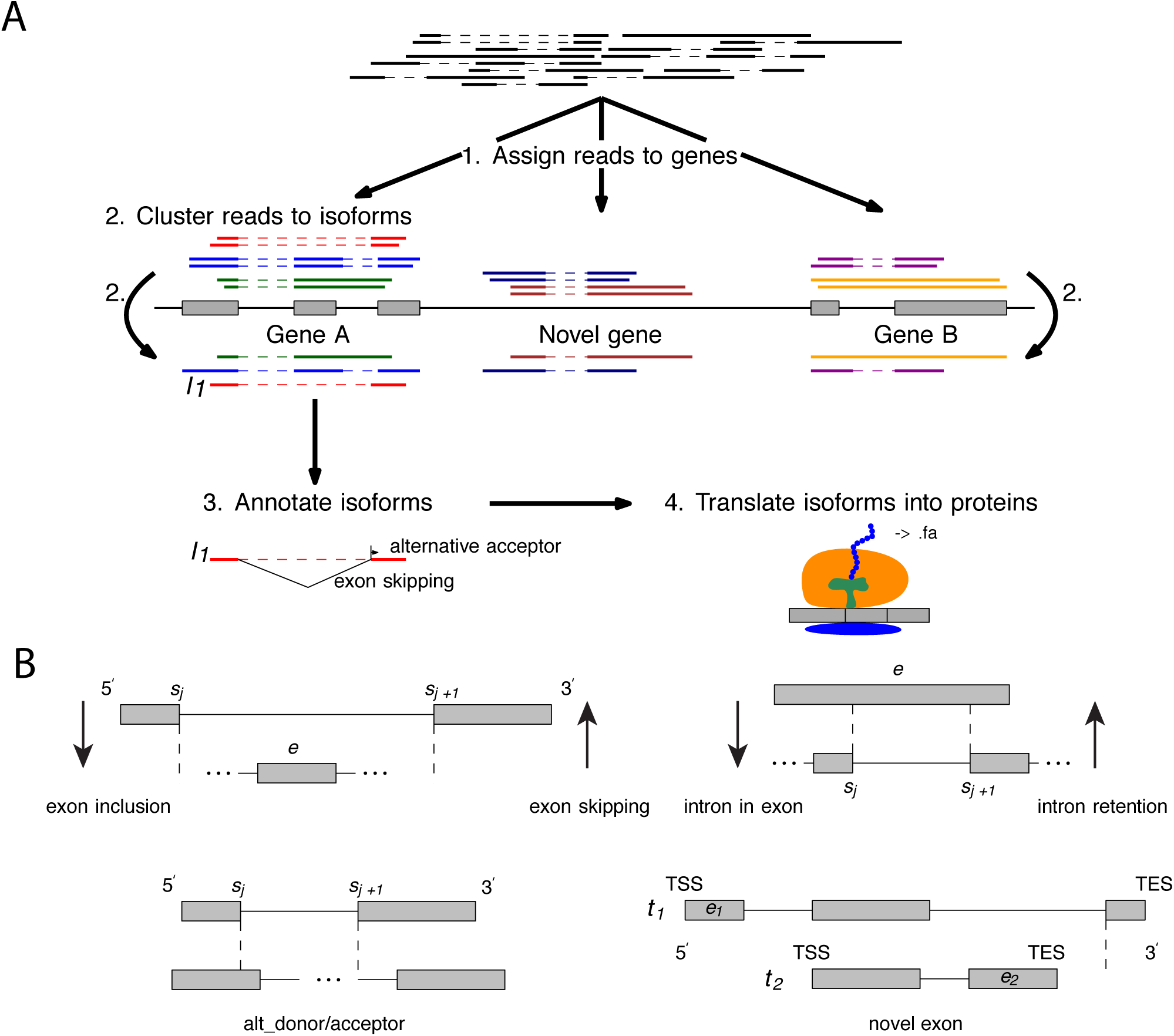
Summary of the transcriptome analysis using PacBio sequencing. (A) A pipeline of SpliceHunter analysis. (B) 8 AS types are defined and examined in this study. Upper left: Exon *e* is strictly contained in intron (*s*_*j*_, *s*_*j* + 1_) and both primary RNA sequences contain sites *s*_*j*_ and *s*_*j* + 1_ if TSS and TES lie beyond the dashed vertical lines. The skipping and inclusion of exon *e* is a symmetric event with respect to the roles of reference and novel transcript. Upper right: Exon *e* retains intron (*s*_*j*_, *s*_*j* + 1_). Intron retention and intron in exon are symmetric events with respect to the roles of reference and novel transcripts. Lower left: *s*_*j*_ and *s*_*j* + 1_ form an alternative donor/acceptor pair. They are contained in both primary RNA sequences if TSS and TES lie beyond the dashed vertical lines. Lower right: Exons *e*_1_ and *e*_2_ are novel in transcripts *t*_1_ and *t*_2_, respectively. Although *e*_2_ lies within the primary RNA sequence of *t*_1_, it does not constitute an inclusion even according to Definition 2 as long as the TES of *t*_2_ lies to the left of the dashed vertical line.

We consider the following types of AS events (Figure 1B). An *exon skipping* is implied by a novel intron that contains at least one complete exon of the reference transcript. An exon in a novel isoform represents an *exon inclusion* event if it is skipped in the reference transcript. An *intron retention* is defined as an exon in the novel isoform that strictly contains at least one intron in the reference transcript. Conversely, we denote an intron of the novel isoform as *intron in exon* if it is retained as part of an exon of the reference transcript. *Alternative acceptors* and *donors* are the 3’ and 5’ ends of an intron that do not appear in the reference transcript. Finally, *novel exons* are not included (see above) and do not overlap any exon in the reference transcript. Formal definitions of all AS events can be found in Methods.

### Characterization of PacBio reads

In total we obtained 2,266,791 CCS reads and 1,311,840 of those were FL CCS reads (Figure 2A, Table S2). 424,511 Isoseq reads were generated through the Iso-Seq pipeline. The average length of CCS reads was 1285 bp, which was slightly longer than previous reports (Figure 2B) (Tilgner et al. 2014; Sharon et al. 2013; Thomas et al. 2014). The average length of FL CCS reads and Isoseq reads were 1094 bp and 1178 bp, respectively. We also examined the length distribution of CCS, FL CCS and Isoseq reads at each time point and did not observe any significant difference across time points (Figure S2). 6307 *Sch. pombe* genes were recovered from PacBio sequencing, which included about 90% of all annotated genes. Median coverage of FL CCS reads per gene was 71 and many genes had coverage >100, which enabled deep discovery of novel isoforms (Figure 2C). 5050 (98.2%) protein-coding genes were recovered which provided a broad scope to explore the relationship between alternative splicing and translation. Additionally, 1154 (75%) non-coding RNAs and 22 (75.9%) pseudogenes were captured so that different regulatory mechanisms could be explored. 97% of the FL CCS reads were assigned to protein-coding genes which represented 80% of all captured genes, indicating higher coverage for protein-coding genes (Figure 2D). Annotated ncRNA genes, which represented 18.3% of captured genes, only accounted for 2.76% of reads and pseudogenes had 0.118% of reads assigned. Interestingly, RNA molecules corresponding to protein-coding genes decreased in the middle of meiosis and increased at the late stage while ncRNA exhibited an opposite pattern (Figure 2E). This may imply anti-correlation mechanisms between certain classes of protein-coding and ncRNA genes. On the other hand, molecules corresponding to pseudogenes monotonically increased during meiosis, suggesting that certain pseudogenes may be up-regulated in meiosis and/or functionally related to meiosis. We further explored the complexity of PacBio reads by examining the number of introns in FL CCS reads. Although the majority of reads contained only 0 or 1 intron, there were still 203,815 reads with multiple introns, together providing substantial data to explore alternative splicing. Lastly, we explored the distances from the 5’ or 3’ end of reads to the corresponding gene transcription start site (TSS) or transcription end site (TES). We focused on reads which extended beyond the annotated TSS or TES sites because the SMARTer cDNA preparation method does not select for 7-methylguanosine (7mG) which marks intact 5’ end mRNA molecules. On average, reads extend 232 bp upstream of 5’ ends (Figure 2G) and 188 bp downstream of 3’ ends (Figure 2H) of annotated genes, suggesting alternative 5’ and 3’ UTRs.

**Figure 2.**
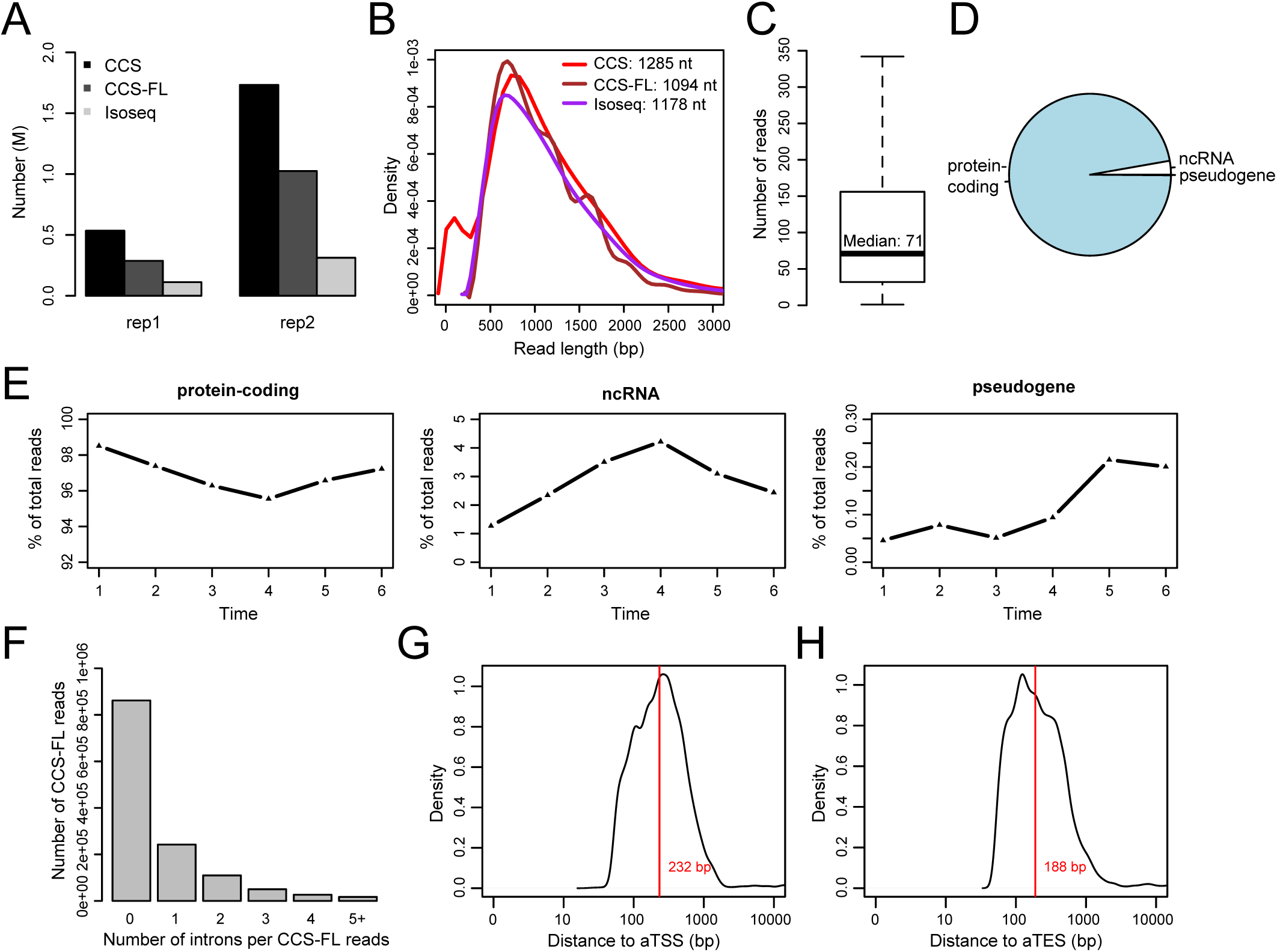
General properties of PacBio reads. (A) Numbers of CCS reads, full-length CCS reads (CCS-FL) and Isoseq reads from two replicates. The second batch has 4x size of the first batch. (B) Length distribution of CCS, CCS-FL and Isoseq reads. (C) Box plot showing the numbers of FL CCS reads for each gene. (D) Pie plot showing the percentage of FL CCS reads corresponding to protein-coding genes, non-coding RNA genes and pseudogenes. (E) Overall trends of FL CCS reads matching protein-coding genes, non-coding RNA genes and pseudogenes. (F) Counts of FL CCS reads with different number of introns. (G) Distribution of distances between the 5’ end of reads to the annotated TSS sites (aTSSs). Reads only outside the annotated TSS sites were counted. (H) Distribution of distances between the 3’ end of reads to the annotated TES sites (aTESs). Reads only outside the annotated TES sites were counted.

### Various types of alternative splicing detected in *Sch. pombe* meiosis

Next, we explored alternative splicing (AS) isoforms in *Sch. pombe*. Although multiple studies have revealed different AS events, this is to our knowledge the first study to explore AS at the isoform level. Not surprisingly, we unveiled complex cases of alternative splicing. For example, we observed 68 distinct polyadenylated mRNA isoforms for *SPAC12B10.05*, containing different types of AS events either in the same or different isoforms (Figure 3A). The capacity to detect multiple AS events in the same molecule by PacBio sequencing avoids underestimating isoform complexity.

**Figure 3.**
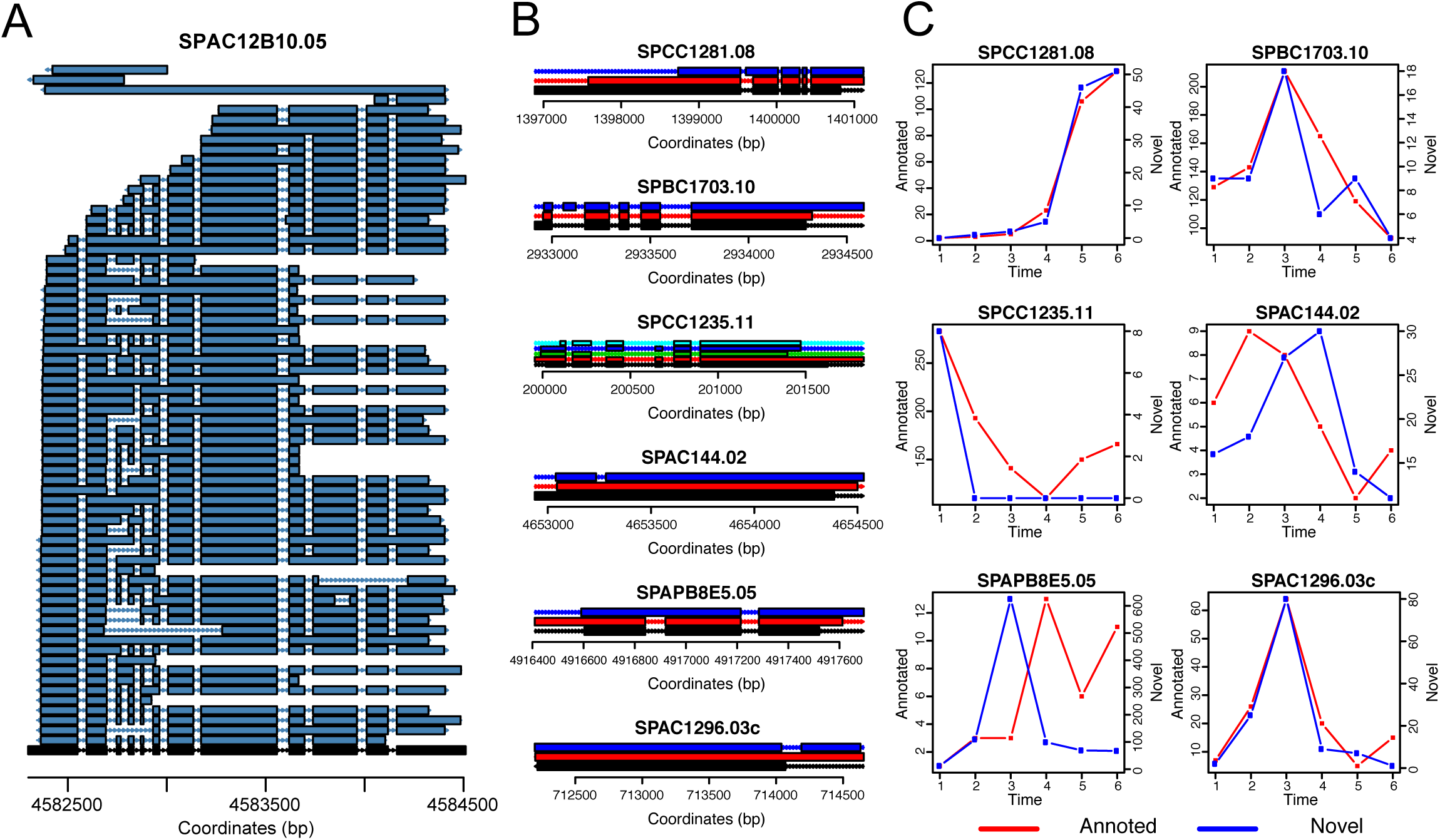
Examples of alternative splicing during *Sch. pombe* meiosis. (A) 68 different isoforms were detected for *SPAC12B10.05*. Black represents the annotation and dark blue represents detected isoforms. (B) 6 different types of alternative splicing structures. *SPCC1281.08* (novel splicing donor or acceptor); *SPBC1703.10* (exon inclusion); *SPCC1235.11* (exon skipping); *SPAC144.02* (intron in exon); *SPAPB8E5.05* (intron retention); *SPAC1296.03c* (novel exon). (C) Dynamics of annotated isoform and novel isoform for each of the 6 examples shown in (B). Red lines represent numbers of FL CCS reads for annotated isoforms and blue lines represent numbers of FL CCS reads for novel isoforms during meiosis.

Alternative splicing comes in many potential forms and guises, and we realized that a formal nomenclature/classification scheme was needed to discuss the many events we observed unambiguously. Thus we developed a rigorous classification scheme and nomenclature that embraces commonly known instances such as exon skipping, intron inclusion, etc. Based on the definitions implemented in our algorithm (see Methods), we observed examples for all types of AS in *Sch. pombe*: an alternative splicing acceptor in *SPCC1281.08*, an exon inclusion in *SPBC1703.10*, multiple exon skipping events in *SPCC1235.11*, an intron in exon in *SPAC144.02*, an intron retention in *SPAPB8E5.05*, and a novel exon in *SPAC1296.03c* (Figure 3B). Interestingly, some of the novel isoforms had comparable (i.e. *SPAC1296.03c*) or even more numerous supporting reads than the corresponding annotated isoforms (i.e. *SPAC144.02* and *SPAPB8E5.05*) (Figure 3C). This implies that these isoforms were unlikely to represent rare erroneous byproducts that escaped mRNA surveillance. Although many novel isoforms showed splicing patterns well-correlated with the corresponding annotated isoforms, some novel isoforms exhibited distinct temporal patterns compared to the corresponding annotated isoforms (Figure 3C), suggesting that the novel isoforms could be temporally differentially regulated. More examples of this are shown in Figure S3.

### Landscape of alternative splicing

After examining individual examples, we next sought to explore the landscape of AS and isoforms (Table S1). The dominant type of AS event observed in *Sch. pombe* was intron retention, probably caused by bypass of individual splicing events (Figure 4A, Table S3). The next most dominant type was intron in exon, which can be considered the opposite of intron retention. Other types of AS events were observed but less abundant, including novel splicing sites, novel exon and exon skipping/inclusion. Next, we asked how many genes have multiple isoforms. Interestingly, only ~1300 genes have a single detected isoform in this study and 1584 genes have two isoforms setting a minimum threshold for consideration of 1 FL CCS reads per isoform (Figure 4B). More than 3000 genes have > 2 isoforms, suggesting pervasive complexity of the *Sch. pombe* transcriptome and potential AS-mediated regulation. Isoform counts per gene were further analyzed by increasing the minimum number of FL CCS reads required to support an isoform (Table S4). Additionally, for the first time we are able to distinguish isoforms with single AS events (9860 or 75%) and mRNA isoforms with multiple AS events (25%) (Figure 4C). Although AS was prevalent in *Sch. pombe*, the annotated isoform was the dominant form in the majority of genes (3614 genes had >90% of reads assigned to the annotated isoform) (Table S5). However, there were also 717 genes with the annotated isoforms accounting for less than the sum of the number of reads supporting alternative isoforms (Figure 4D). In general, FL CCS reads matching annotated isoforms were ~8 fold more abundant than the reads matching novel isoforms. Besides typical AS isoforms, we also discovered mRNAs apparently encoding 887 new TUs (Table S1) and ~4000 antisense isoforms (Table S6) based on a minimum read depth of 1. 722 of the new TUs had no intron with a maximum of 262 supporting reads and 165 new TUs had introns with a maximum of 61 supporting reads. 568 of 635 annotated antisense RNA genes were detected from our data and we additionally detected antisense isoforms targeting 3589 other genes, consistent with a previous study showing the prevalence of antisense meiotic transcripts (Ni et al. 2010). However, the number of reads supporting novel antisense isoforms were generally substantially lower than the number of reads matching annotated antisense isoforms (Figure 4E). Therefore, there was potentially a mix of real antisense isoforms and “transcriptional noise”. Furthermore, we examined dinucleotides at the novel splicing sites. 99.94% of the dinucleotides at annotated splicing sites are GT-AG, with only 3 exceptions. However, GT-AG only appeared at 64.29% of the novel splicing sites with all types of variations after excluding alignment errors, implying that many of these novel introns might be aberrantly spliced (Table S7). Pairs of dinucleotides with small “hamming distances” to GT-AG occurred a bit more often, such as 79 pairs of GT-TG, 68 pairs of GT-TG etc.

**Figure 4.**
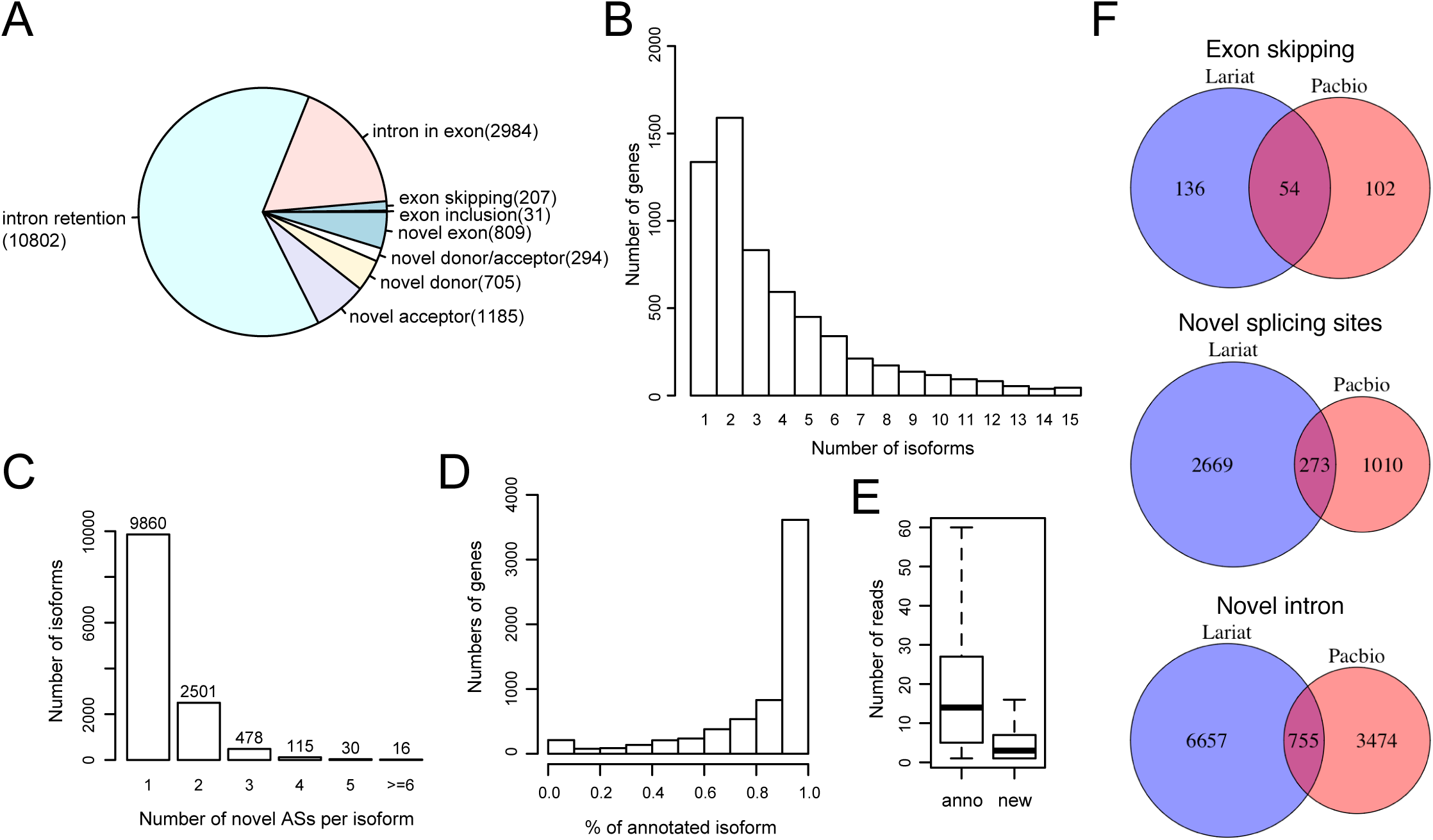
Landscape of alternative splicing events during *Sch. pombe* meiosis. (A) Pie chart showing the fraction of different types of alternative splicing events in *Sch. pombe*. Numbers of alternative splicing for each type are listed after the names. (B) Count of genes with different number of isoforms. (C) Isoforms broken up by the number of novel alternative splicing events discovered in individual isoforms. (D) Genes broken up by the percentage of FL CCS reads corresponding to the annotated isoforms. (E) Box plot showing the numbers of FL CCS reads supporting annotated and novel anti-sense transcripts. (F) Venn graphs showing the comparison between AS events detected by PacBio sequencing and lariat sequencing.

Interestingly, we discovered some read-through transcripts that spanned two consecutive genes on the same strand, 60 of which contained annotated introns from both genes (Table S1, S8). Among read-through transcripts with introns, 15 had supporting reads more abundant than at least one of the individual transcripts, suggesting that these were less likely to represent chimeric reads. This is consistent with a previous finding of widespread “polycistronic” transcripts in fungi (Gordon et al. 2015). Figure S4 shows 4 examples of read-through transcripts. Read-through transcripts for *SPBC887.05c-SPBC887.06c, SPAPB1A10.03-SPAPB1A10.16* and *SPAC19D5.05c-SPAC19D5.11c* covered every exon of individual genes but *SPAC16E8.02-SPAC16E8.03* lacked the last, noncoding exon of *SPAC16E8.02*, which theoretically leads to an in-frame fusion of the two ORFs (Figure S4D). *SPAC16E8.02* encodes a DUF962 family protein and *SPAC16E8.03* (*gna1*) encodes a glucosamine-phosphate N-acetyltransferase. Interestingly, both genes are essential for spore germination (Hayles et al. 2013). Importantly, the read-through transcript was highly induced at the late stage of meiosis but individual transcripts were low and static, suggesting that the read-through transcript encodes a fused protein essential for spore germination. Another interesting example is *SPAC19D5.05c-SPAC19D5.11c*, which exhibited the opposite temporal pattern compared to *SPAC19D5.05c*(*imp3*). Imp3 is a predicted U3 snoRNP-associated protein and it is required for both vegetative growth and spore germination (Hayles et al. 2013; Kim et al. 2010). Interestingly, the transcript of *imp3* decreased from mitosis to meiosis while the read-through isoform increased in the middle of meiosis (Figure S4C), implying a possible switch of the transcriptional program for imp3 between mitosis and meiosis. The two reading frames are not disrupted in this read-through isoform.

Different methods have been applied in *Sch. pombe* to identify alternative splicing, including short-read RNA-seq, ribosomal profiling and lariat sequencing (Duncan and Mata 2014; Bitton et al. 2015; Awan et al. 2013; Stepankiw et al. 2015). Here we compared our results with corresponding events predicted from lariat sequencing and ribosomal profiling. Ribosomal profiling was performed in meiosis too (Duncan and Mata 2014)and excepting novel start and stop codons, which were not predicted by our study, the remaining 8 new isoforms detected were all validated in our study. Lariat sequencing was done in a *dbr1* mutant strain from log phase, diauxic shift and heat shock conditions (Stepankiw et al. 2015). The occurrences of three types of AS events were compared, including exon skipping, novel splicing sites and novel intron. Surprisingly, all the three types of AS events from the two datasets had little overlap (Figure 4F), probably because the conditions from the two studies were distinct and the techniques captured different RNA molecules. The comparison suggests that the complexity of AS is even higher.

### Dynamics of AS during meiosis

When we examined AS events, we noticed that many alternative isoforms exhibited unexpected temporal patterns compared annotated isoforms. We next sought to explore the dynamics of AS isoforms. We first summarized the general dynamic patterns of different types of AS events by examining the number of isoforms and number of reads supporting each type of AS. Most types of AS increased in meiosis (Figure 5A), which was likely to be at least partially related to decreased RNA surveillance. However, exon-skipping events decreased at early stages of meiosis and increased at the late stage; intron retention was relatively unchanged. The dynamics of exon skipping suggest a condition-driven alternative usage of different exons between mitosis and meiosis.

**Figure 5.**
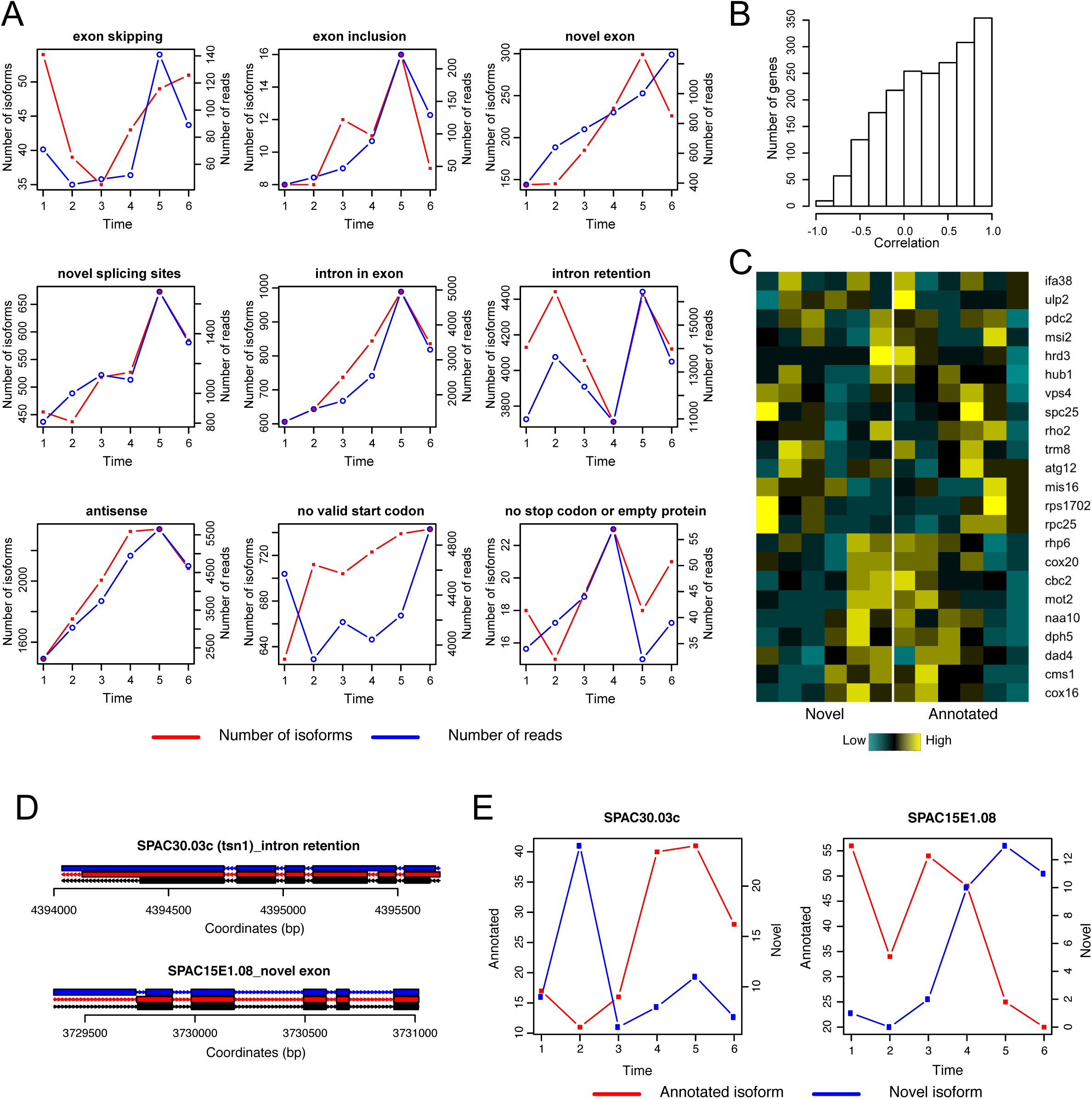
Dynamics of alternative splicing during meiosis. (A) Trends of different types of alternative splicing during meiosis. Red lines represent numbers of isoforms and blue lines represent numbers of FL CCS reads. (B) Histogram showing the correlation between annotated and alternative isoforms during meiosis. (C) Heat map showing the temporal pattern of novel and annotated isoforms. Genes with correlation smaller than ‐0.5, FL CCS read count > 100 and percentage of annotated isoform < 90% were chosen for hierarchical clustering. (D-E) Examples of anti-correlated isoforms in meiosis. (D) shows the exon-intron structures of the examples. (E) shows the temporal patterns of the annotated and alternative isoforms.

Next, we examined isoform dynamics at the gene level. We hypothesized that if AS is functionally associated with different conditions, the two isoforms for the same gene are more likely to be un-correlated or anti-correlated. Therefore, Pearson’s correlation coefficients were calculated between the number of reads supporting annotated and alternative isoforms for genes with comparable abundance of each isoform and coefficients were displayed in a histogram (Figure 5B, Table S9). A left skewed shape suggests that the majority of alternative isoforms was correlated in abundance with annotated isoforms. However, there were still 580 genes with anti-correlated isoforms and 109 genes with correlation <= –0.5. To explore the anti-correlated isoforms, we selected 28 genes with Pearson correlation <= –0.5 that were supported by more than 100 FL CCS reads for further analysis. 5 of them were predicted genes, which we excluded. The temporal patterns of novel and annotated isoforms of the remaining 23 genes were compared in a heat map (Figure 5C). Some genes had alternative isoforms expressed in mitosis and annotated isoforms expressed in meiosis whereas a second cluster of genes had annotated isoforms expressed in mitosis and switched to alternative isoforms in late meiosis.

*SPAC30.03c* (*tsn1*) encodes a translin protein (Laufman et al. 2005)and it is up regulated in the middle and late meiosis (Martín-Castellanos et al. 2005; Mata et al. 2002). Interestingly, besides an increment of annotated isoform expression in middle to late meiosis, we also observed an intron retention isoform expressed in early meiosis (Figure 5D, E). A 48 bp intron was retained, which kept the predicted translation in frame and terminated at the annotated stop codon. *SPAC15E1.08* (*naa10*) encodes a N-acetyltransferase catalytic subunit, which is essential for survival, and mutations in the *S.cerevisiae* homolog *ARD1* cause sporulation defects (Mullen et al. 1989) (Mullen JR 1989). The abundance of the novel isoform which has a novel exon and an intron upstream of the annotated start codon, increased in late meiosis. It is possible that the new 5’ UTR is involved in the regulation or provides new start sites. A few examples were further examined in Figure S5. Two novel introns were found in the coding region of *SPCC162.04c* (*wtf13*), which did not disrupt the reading frame and retained the annotated stop codon. This isoform should be the dominant isoform and its expression increased in late meiosis. We also observed a typical class of anticorrelated isoforms, antisense RNA isoforms for *SPBC23G7.07c* (*cms1*) and *SPBC28F2.08c* (*hrd3*). The antisense isoforms increased in late meiosis, when the annotated isoforms decreased, suggesting an inhibitory effect of antisense isoforms. Additionally, we observed a few more examples of intron retention events with anti-correlated dynamics, such as *SPAC16C9.04c* (*mot2*), *SPBC3B9.22c* (*dad4*) and *SPCC576.14* (*dph5*). However, these intron retentions lead to early termination of predicted translation by introducing stop codons in the retained introns.

### Prediction of novel proteins

One major role of alternative splicing is to generate different functional proteins from the same gene. On the contrary, a product of aberrant splicing is not responsible for generating meaningful proteins. Therefore, we performed systematic prediction of protein sequences from all isoforms detected in our study (Table S10). Translation prediction always started from the annotated start codon if it is available and stopped at the first stop codon. By this definition, 19081 isoforms were identified as translatable and the remaining 9298 isoforms could not generate valid proteins (Figure 6A). 15102 different protein sequences were predicted from 5029 genes. Among these genes, 2933 genes were predicted to generate exclusively annotated proteins and 2080 genes were predicted to generate both annotated and novel proteins. Interestingly, 16 genes were assumed to generate novel proteins only (Table S11). Some of them are likely to be misannotated.

**Figure 6.**
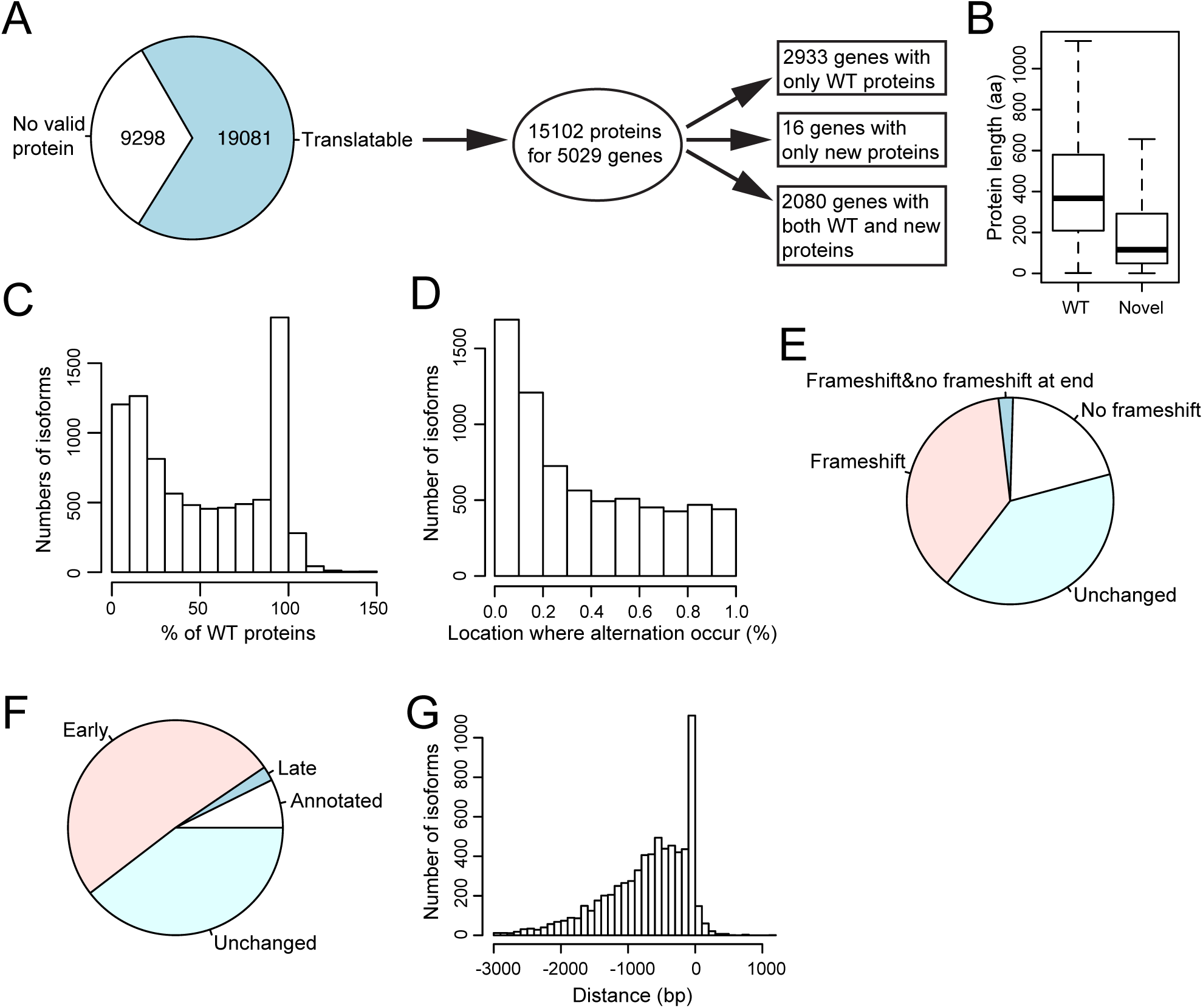
Predicting translational products from detected isoforms. (A) Summary of protein prediction. The left pie plot shows the number of isoforms generating proteins. 15102 proteins were predicted fro 5029 genes and these genes were further broken up by the status of annotated and new proteins. (B) Box plot indicating the length of annotated and novel proteins. (C) For isoforms generating novel proteins, a histogram shows the ratio of novel proteins relative to corresponding annotated proteins. (D) A histogram showing where alternation occurs in the new protein relative to the corresponding annotated protein. (E) Fractions of translatable isoforms with novel AS events which contain frameshift or not. Unchanged were isoforms with same proteins as annotation. (F) Fractions of translatable isoforms with novel AS events which have early, late or annotated stop codons. (G) A histogram showing the distances between annotated and novel stop codons for isoforms with novel predicted proteins.

A predicted novel protein may not actually be produced. We sought to theoretically address this question by examining important protein sequence properties, including its length, relative position of protein sequence alteration caused by AS, how it affects the reading frame and where the translation of novel proteins terminate compared to that of annotated proteins. In general, novel protein sequences were shorter than annotated protein sequences (Figure 6B). We observed an enrichment of isoforms which generated novel proteins with similar length to annotated proteins and we also observed an enrichment of isoforms which generated novel proteins with <20% of annotated protein length (Figure 6C). The first class of isoforms might be actually translated but the second class is considered unlikely to produce functional proteins. Furthermore, protein alteration was more likely to occur at the beginning of translation (Figure 6D). Overall, 39.6% of translatable isoforms with AS events encoded the same proteins as annotated isoforms. More than half of the remaining isoforms which generate novel proteins are partially not in the same reading frame as annotated isoforms (Figure 6E). Importantly, ~33.76% of isoforms producing novel proteins were in the same reading frame as annotated isoforms and another ~3.75% of isoforms had alterations that shifted the reading frame followed by a second alteration that restored the frame. Furthermore, 7.3% of AS isoforms had translation ending at the annotated stop codons (Figure 6F). Only for 2.17% of AS isoforms, translation ended downstream of, but close to, annotated stop codons (Figure 6G). Most isoforms with altered protein sequences (50.9%) showed early translation termination with varying distances between annotated and novel stop codons.

In addition, we attempted to predict protein coding regions for the 887 new TUs, some of which are potential mRNAs for previously unknown genes. The protein length was generally very short 12 with a median length of around 30 aa; the longest was >200 aa (Figure S6). The majority of novel TUs had no ORF longer than 25% of their full-length sequence. Therefore, it is likely that the majority of these novel transcripts represent ncRNAs. Previous work has hypothesized that some or most ncRNAs represent “transcriptional noise” (Costa 2007; Hüttenhofer et al. 2005) but may also represent fodder for the birth of new genes (Carvunis et al. 2012). This is supported by the observation that only 13% of the novel TUs identified had ribosome signals in a previous profiling study (Duncan and Mata 2014).

### Protein sequence conservation and secondary structure

Following the protein prediction analysis, we next sought to check the quality of these novel protein sequences by evaluating their conservation and secondary structure formation. Two major classes of novel protein sequences were examined. Hypothetic proteins predicted from novel TUs and novel C terminal amino acid sequences starting at the first alteration in AS isoforms. Conservation of novel sequences was assessed by searching for sequence-similar proteins in fission yeasts, fungi and eukaryotes using BLAST (Gish and States 1993). Secondary and tertiary structures and properties were predicted by RaptorX (Källberg et al. 2012). For 59 novel TUs we found evidence for homology in other species with less than 100 matching aa (Figure S7D). The majority of them had homologs in other related fission yeasts without significant preference (Figure 7A). This supports the idea that the majority of new TUs are not being translated but others may be translationally active. For example, the predicted protein of NG_629 highly resembles a membrane transporter such as allantoate permease in other fission yeasts and it could form a structure with 64% α-helix (Figure S7A). NG_634 is predicted to encode a protein similar to RecQ type DNA helicase and the predicted structure contains 34% α-helix, 25% β-sheet and 40% coil (Figure S7B). The longest protein sequence predicted from a novel TU (NG_653) is similar to retrotransposons in different fission yeasts (Figure S7C). Additionally, we examined conservation of C terminus novel amino acid sequences from 614 AS isoforms which have >2 CCS FL supporting reads and are at least 19 aa long. 183 AS isoforms have homologs in other species, predominantly in fission yeasts (Figure 7B).

**Figure 7.**
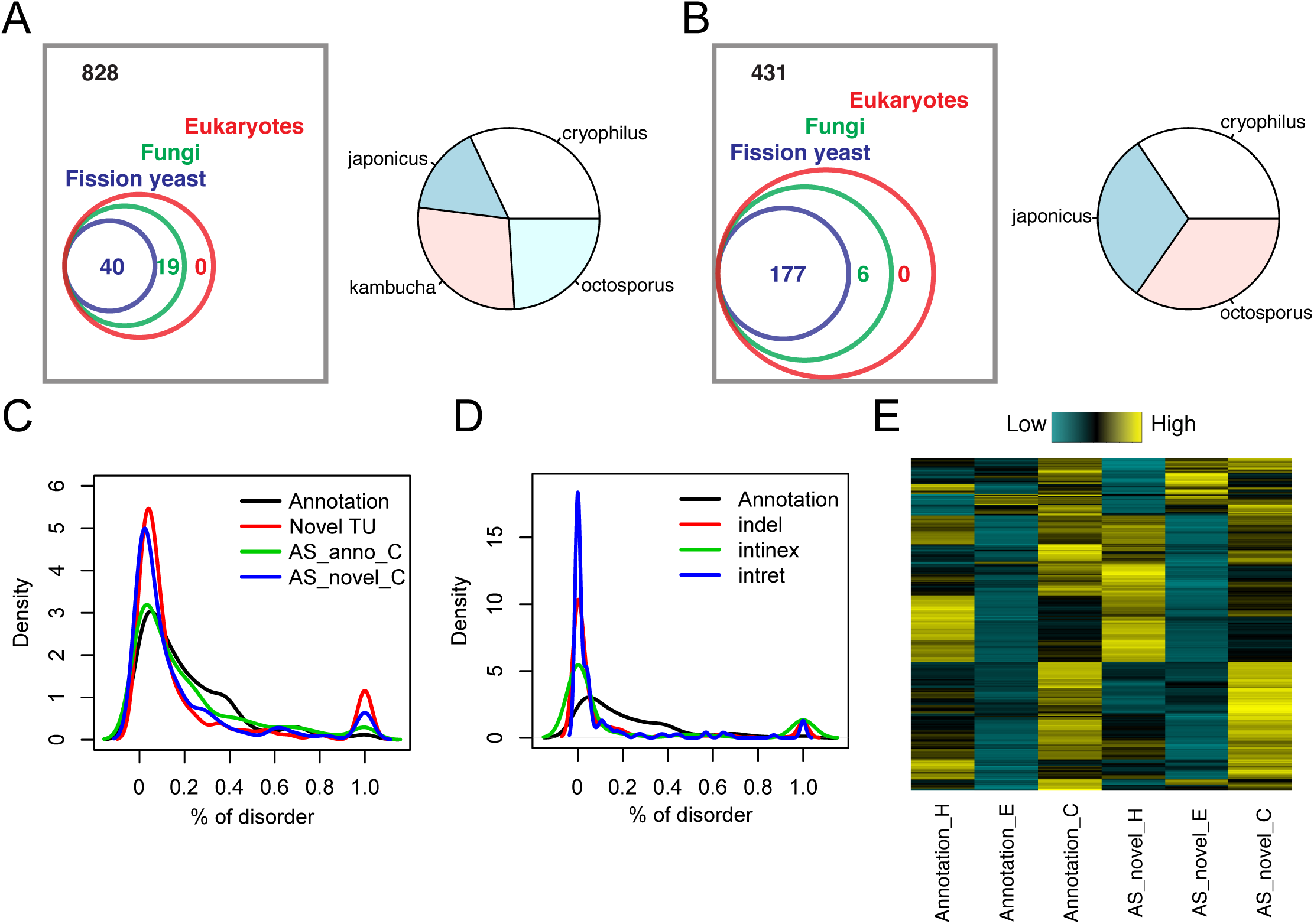
Conservation and secondary structure analysis. (A) Left panel shows the protein blast results of the novel TUs. Numbers of novel TUs with alignment hits in different taxa are shown. Right panel shows the proportion of hits in different fission yeasts. (B) Left panel shows the protein blast results using only the modified part of proteins from AS isoforms. Numbers of AS isoforms with alignment hits in different taxa are shown. Right panel shows the proportion of hits in different fission yeasts. (C) Distribution of percent of disorder predicted by RaptorX (Källberg et al. 2012). 4 classes of protein sequences were examined: full length annotated proteins, longest proteins predicted from novel TUs, C terminus annotated or amino acid sequence starting at the first alteration in AS isoforms. (D) Distributions of percent of disorder for annotated proteins, amino acids affected by insertions and deletions (indel), intron retentions (intret) and introns in exons (intinex) from AS isoforms. (E) Proportions of α-helix (H), β-sheet (E) and coil (C) predicted from C terminus annotated or amino acid sequence starting at first alteration in AS isoforms.

Next, we systematically evaluated secondary structure formation of novel proteins. The majority of proteins from novel TUs shows very low levels of disorder and reflects the presence of alpha helix or beta sheet propensity. However, a minor mode indicates novel TUs with complete disorder, which is distinct from annotated proteins (Figure 7C). A similar pattern was observed in novel C terminus proteins from AS isoforms versus annotated C terminus proteins. We further explored this pattern of disordered residues among inserted (by intron retentions) and deleted (by introns in exons) amino acid sequences. We consistently observed a prime enrichment of low disorder and a secondary enrichment of complete disorder (Figure 7D). Then we asked how alternative C terminal amino acids in AS isoforms affect secondary structures. Figure 7E and S7E suggest that the majority of altered C termini maintain the proportion of α-helix (H), β-sheet (E) and coil (C). Various classes of modified amino acid sequences show a similar composition of secondary structure elements (Figure S7F).

## Discussion

In summary, we systematically characterized the transcriptome diversity and dynamics at the isoform level during fission yeast meiosis. We identified thousands of novel isoforms and AS events, including unexpected findings such as novel TUs and read-through transcripts. We further carefully evaluated the functional relevance of AS isoforms by scrutinizing three factors, the abundance of AS isoforms, the differential regulation of annotated and AS isoforms during meiosis and the translational capacity and quality of AS isoforms. The majority of AS isoforms show similar temporal patterns compared to the annotated isoforms and the majority of AS isoforms also have shifted reading frames and short predicted protein sequences. However, a few genes show anti-correlated abundance of annotated versus AS isoforms and these AS isoforms have similar length of translatable sequences, suggesting the possibility of AS-mediated regulation of gene activity in the unicellular organism. We developed SpliceHunter, a novel computational tool for the discovery of AS events and isoform-based sequence analysis from PacBio long sequencing reads. SpliceHunter is available at https://bitbucket.org/canzar/splicehunter and can process single or multiple samples (e.g. time series). Our results show that the analysis of isoform dynamics is feasible using PacBio sequencing given a small transcriptome and smooth trends.

How complex is the “functional” transcriptome in *Sch. pombe?* In agreement with previous studies (Bitton et al. 2015; Awan et al. 2013; Stepankiw et al. 2015), our study shows that AS events are widespread in this unicellular organism. Could it be even more complex than this? Although we captured all AS events identified from prior ribosomal profiling data in meiosis, our AS events had a relatively small overlap with the AS events identified by lariat sequencing performed in log phase, diauxic shift and heat stress (Figure 4F). This observation supports two possible conclusions: first, it seems likely that certain AS events could be distinctly regulated under different conditions, which would suggest that some AS events occur condition-specific; second, AS events shared across conditions may represent instances of “constitutive AS” and are more likely to be functionally relevant. They occurred consistently under different conditions and were detected using different techniques and thus seem unlikely to be the result of random splicing errors. Additionally, we focused on isoforms with alternative internal structures, given the limitation of our cDNA library preparation strategy in capturing 5’ intact mRNAs. The diversity of AS increased substantially when we considered isoforms with precise 5’ and 3’ ends different from the annotated ones. Given the differences in AS events between the two studies, a substantially larger diversity of isoforms might be revealed in this unicellular organism when examining additional extrinsic factors like different treatments and environments. On the other hand, in plants and animals AS events can vary across different tissues or organs. Therefore, it might be interesting to explore AS events in fission yeast with varied “developmental” status besides mitosis and meiosis, such as AS in young vs. aged cells or mother vs. daughter cells.

Is an isoform truly functional or just a result of splicing errors? This is a critically important question transcending the overall complexity of detected isoforms. Multiple studies have suggested that a majority of AS events is aberrant and not functional despite their prevalence (Bitton et al. 2015; Stepankiw et al. 2015). This hypothesis is supported by (i) the small number of reads supporting AS events compared to the number of reads supporting annotated splicing, (ii) an increased prevalence of AS events in mutants of RNA surveillance, and (iii) AS events failing to lead to meaningful translational products and ribosomes that are unlikely to be bound (Bitton et al. 2015). Our study is in line with this conclusion from the isoform perspective. Most genes have >90% reads supporting annotated isoform (Figure 4D); the majority of AS isoforms lead to frameshifts and shorter than annotated protein sequences (Figure 6C,D,E), although we describe several interesting exceptions. We additionally show that a majority of genes have correlated abundance between annotated and AS isoform during meiosis, suggesting a lack of regulated AS from the temporal perspective for these genes. Combining all these characteristics, we suspect that a large portion of the novel isoforms is not translationally active. However, the possibility of functional AS is not excluded and could be condition-specific. AS in the *alp41* and *qcr10* genes were specifically induced by heat or cold shock (Awan et al. 2013).

On the other hand, we find many excellent candidate novel transcripts that are likely to be functional. For example, we identified a few hundred genes that have less reads assigned to the annotated isoforms than to the AS isoforms as described here. Moreover, certain AS isoforms were dynamically regulated during meiosis, and some of them even exhibited anti-correlated expression patterns compared to corresponding annotated isoforms (Figure S5). These seem quite likely to be functionally relevant. Although many of them were still not likely to be translated, we suspect that AS could explain the regulation of some of these genes in meiosis. Furthermore, the combination of conservation analysis and secondary structure modeling indicates that the predicted proteins are potentially able to form stable structures. We were able to infer the functions of some novel proteins through their homologs in other related organisms. Future studies are needed to confirm the existence of predicted proteins and diagnose potential physiological functions. Additionally, AS isoforms could play regulatory, noncoding roles, similar to what was observed for some of the anti-sense isoforms. Finally, the fact that many novel TUs encode protein isoforms not found in closely related species could represent simply a low level of “background noise” arising from aberrant splicing, but it could also represent a mechanism for the evolution of new gene function; some subset of these reading frames may represent species-specific new gene isoform birth (Carvunis et al. 2012).

Overall, these data indicate that fission yeast has a large repertoire of AS events. Although the majority may not function to produce altered proteins, some of these may form a kind of substrate for the evolution of condition-specific isoforms. The remainder could also serve as a resource for neofunctionalization or new isoform birth. Finally, this dataset will be a valuable resource for studying AS as a regulatory mechanism for a given gene of interest in this widely used model organism.

## Methods

### General methods

Standard methods were used for growth and meiosis of *Sch. pombe*. Briefly, SP2910 (*mat1-PA 17/mat1-M-smt-o his2/+ leu1-32/+ ura4/+ ade6-M210/ade6-M216*) was grown in 1ml YES medium overnight and diluted in EMM culture at an OD of 0.1. Cells were harvested and washed when they entered into the log phase (~OD 0.6) and resuspended in EMM without NH_4_Cl (EMMN). 10 OD of cells were collected from 0 to 10 hours after resuspending in EMM-N medium every 2 hours. Total RNA was extracted using the RNeasy Mini kit (Qiagen, 74106) and polyA selection was performed using the Poly(A)Purist MAG kit (Invitrogen AM1922). cDNA libraries were synthesized using the SMARTer PCR cDNA synthesis kit (Clontech, 634926). SMRT libraries were generated in the Johns Hopkins Deep Sequencing and Microarray core facility using the Pacific Biosciences’s template prep kit. 5 SMRT cells were used for each time point.

### Iso-Seq analysis and read alignment

Iso-Seq analysis was performed on the SMRT portal website. Reads from different replicates and time points were processed independently. Standard parameters were used to cluster and polish reads. We used the CCS, full length CCS and Isoseq reads for downstream analysis. The number of reads and and their length were directly obtained from read sequence files in fasta format. Isoseq reads were aligned to the v2.29 *Sch. pombe* genome (Wood et al. 2012) using GMAP (Wu and Watanabe 2005). The alignments were provided to the SpliceHunter pipeline which initially clusters isoseq reads into isoforms. For each isoform, the coordinates and composition of exons are compared to annotated exon-intron structures to infer the types of AS events. Details of the SpliceHunter pipeline are described below. R was used to generate the plots.

### Comparison of AS events

Alternative exon/intron positions from the ribosomal profiling study were manually compared to our results since only 13 genes were reported to have alternative coordinates of exon intron junctions. AS events from the lariat sequencing results were compared by firstly generating the pairs of 5’ and 3’ splicing sites from both studies. Combinations of coordinates were compared between the two studies to infer the overlap. Besides exact matches, we also relaxed the exon/intron boundaries by +/-1, 2, 3 bp and searched for overlapping pairs. We did not observe large differences by relaxing the boundaries. Results based on perfect matches were displayed in Venn diagrams.

### Conservation analysis

We performed a BLAST search of novel protein sequences against taxa databases of fission yeast, fungi and eukaryotes. Alignments with alignment length >= 30 aa and E value <= 0.05 for fission yeast and fungi or E value <= 0.001 for eukaryotes were considered as hits. For novel TUs, we extended our search for homologs by running tblastn and tblastx (Altschul et al. 1997), while changing the minimum alignment length to 50 bp. In this analysis, we only considered amino acid sequences from AS isoforms that had >2 supporting FL CCS reads and where the sequence starting at the first AS event was >=19 amino acids long. Secondary structure was predicted using the RaptorX web portal.

### Alternative Splicing Event Definition

We introduce a rigorous classification scheme and nomenclature of alternative splicing events that enables an unambiguous discussion of observed splicing patterns. We represent a transcript *t* by its sequence of splice sites 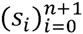, ordered from 5’ to 3’. Sites *s*_0_ and *s*_*n* + 1_ represent transcription start and end site, respectively, and (*s*_2*j* + 1_, *s*_2*j* + 2_) the introns, for 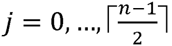. The implied exons are (*s*_2*j*_, *s*_2*j* + 1_) for 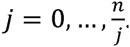. We assign alternative splicing events of a transcript 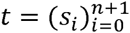 with respect to a reference transcript 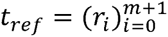 to one of the following categories (see Figure 1B).

#### Definition 1 (Exon skipping)

A novel intron (*s*_*j*_,*s*_*j*+1_) in *t* implies an *exon skipping* event if the interval [*s*_*j*_, *s*_*j*+1_] strictly contains at least one exon *e* of *t*_*ref*_ and if both *s*_*j*_ and *s*_*j*+1_ are present in the primary RNA sequence of *t*_*ref*_.

#### Definition 2 (Exon inclusion)

An exon *e* in *t* implies an *exon inclusion* event if it is skipped (according to Definition 1) in *t*_*ref*_ with respect to *t*.

#### Definition 3 (Intron retention)

An exon *e* in *t* that strictly contains at least one intron in *t*_*ref*_ is denoted as *intron retention*. Note that by definition the splice sites of the retained intron are present in the primary RNA sequence of *t*.

#### Definition 4 (Intron in exon)

We denote an intron (*s*_*j*_, *s*_*j*+1_) as *intron in exon* if it is retained (according to Definition 3) in an exon of *W*.

#### Definition 5 (Alternative donor)

Splice site *s*_*j*_ of an intron (*s*_*j*_,*s*_*j*+1_) in *t* is an *alternative donor*, if it does not match any splice site in *t*_*ref*_ and if it is present in the primary RNA sequence of *t*_*ref*_.

#### Definition 6 (Alternative acceptor)

Splice site *s*_*j*+1_ of an intron (*s*_*j*_, *s*_*j*+1_) in *t* is an *alternative acceptor*, if it does not match any splice site in *t*_*ref*_ and if it is present in the primary RNA sequence of *t*_*ref*_.

#### Definition 7 (Alternative donor/acceptor)

Splice sites *s*_*j*_ and *s*_*j*+1_ of an intron (*s*_*j*_, *s*_*j*+1_) in *t* form an *alternative donor/acceptor* pair, if *s*_*j*_ is an alternative donor (Definition 5) and *s*_*j*+1_ is an alternative acceptor (Defintion 6).

#### Definition 8 (Novel exon)

We define an exon *e* in *t* that does not overlap any exon in *t*_*ref*_ and that is not an exon inclusion (Definition 2) as *novel exon*.

### Alternative Splicing Detection by SpliceHunter

Initially (step 1 in Figure 1A), reads are mapped to a reference genome and assigned to annotated genes (Wood et al. 2012) based on matching introns, matching splice sites, or exonic overlap, in this order. If a read does not span any introns or no splice site matches any annotated splice site, the exonic overlap with annotated genes is considered. If the read alignment overlaps multiple genes, the gene with maximal overlap is chosen, provided that the overlap is significantly larger (default 1.5x) than the second largest overlap. Reads that do not overlap any annotated gene are used to infer *novel TUs*. Ambiguous reads that overlap multiple genes equally well (less than 1.5x difference) were excluded from further analysis.

For each gene and novel TU, all assigned reads are clustered to isoforms by their intron chains and their start and end sites (step 2 in Figure 1A). Single exons reads are grouped by the (potentially empty) sequence of retained annotated introns instead. While intron chains and retained introns are required to match perfectly, start and end sites of reads in the same cluster must lie within a sliding window of adjustable size. The start and end site of the isoform representing such a cluster is defined to be the most 5' start and most 3' end site, respectively, of any read contained in the cluster. For annotated genes, start and end sites of assigned reads that lie close (default 50 bp) to the annotated start or end site, respectively, are snapped to the annotated sites. Single exon reads that were not assigned to an annotated gene are grouped to novel TUs simply based on their overlap.

Third (Figure 1A, step 3), each isoform (sense) is compared to the exon-intron structure of the gene to which it was assigned to detect alternative splicing events (Definitions 1-8). Single-exon isoforms (sense) are tested for intron retention events only. Start and end sites of isoforms that do not show any alternative splicing are annotated as *truncated, novel* (extends beyond the annotated site), or *as annotated*. Antisense transcripts are marked in the output as such. Isoforms whose introns match introns of multiple genes on the same strand are output as *read-through transcripts*. If the matching genes lie on different strands or different chromosomes the isoform is reported as a *fused RNA molecule*.

Finally, the effect of alternative splicing events on the protein sequence is studied (step 4 in Figure 1A). Each (truncated) isoform that has been assigned to an annotated gene is extended on both ends, consistent with the exon-intron structure of the annotated transcript. If the start or end site of the isoform falls into an intron of the annotated transcript, the isoform cannot be extended in 5' or 3' direction, respectively. If the (extended) isoform stretches beyond the position of the annotated start codon, the isoform is translated starting at the annotated start codon. For all isoforms whose structures imply an open reading frame, the corresponding protein sequences are output in fasta format. For all remaining isoforms, the negative translation status is marked accordingly. For novel TUs, the longest ORF is reported.

Furthermore, for each isoform all alternative splicing events are annotated with the number of inserted or deleted amino acids and the resulting shift in reading frame. SpliceHunter aligns novel protein sequences with corresponding annotated sequences and outputs a sequence of strings with pattern Tx_y^^^z. T is either “D” or “I” and indicates whether the alignment gap is a deletion (“D”) or insertion (“I”), x gives the length of the gap, y the position with respect to the reference sequence, and z the frame (0,1,2) following the current gap. At the end of the string, =d+/- denotes the positive or negative distance of the active stop codon from the annotated stop codon.

When provided with samples from multiple replicates across different time points, SpliceHunter pools all samples to cluster reads to isoforms (see second step) but keeps track of the distribution of supporting FL CCS reads across time points (see 8^th^ column in Table S1) or (optionally) across individual samples.

## Data and code access

PacBio sequencing data has been deposited in the Gene Expression Omnibus database under accession number GSE79802.

SpliceHunter is free open-source software released under the GNU GPL license, and has been developed and tested on a Linux x86_64 system. SpliceHunter’s source is available at https://bitbucket.org/canzar/splicehunter.

## Acknowledgements

We thank Haiping Hao for assistance with sequencing, and Alix Kieu and Roberto LIeras, Pacific Biosystems, for in kind contribution of SMRT cells and suggestions for analysis. We thank Ed Miller for providing funding to the High Throughput Biology Center to acquire the PacBio sequencer, and Steven Salzberg for helpful discussions.

## Figure legend

**Figure S1.**
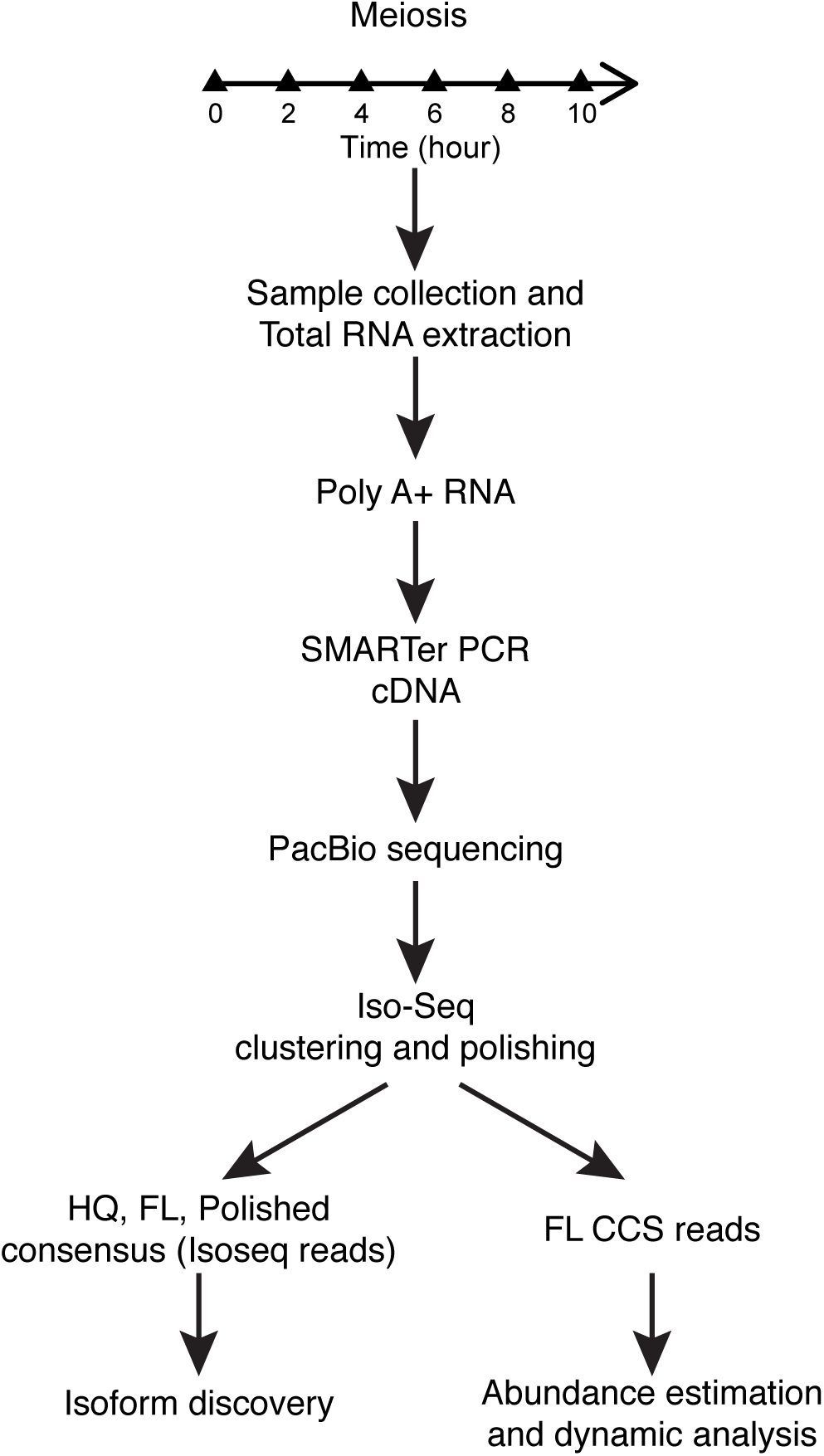
A workflow to characterize the diversity and dynamics of transcriptome in *Sch. pombe* meiosis.

**Figure S2.**
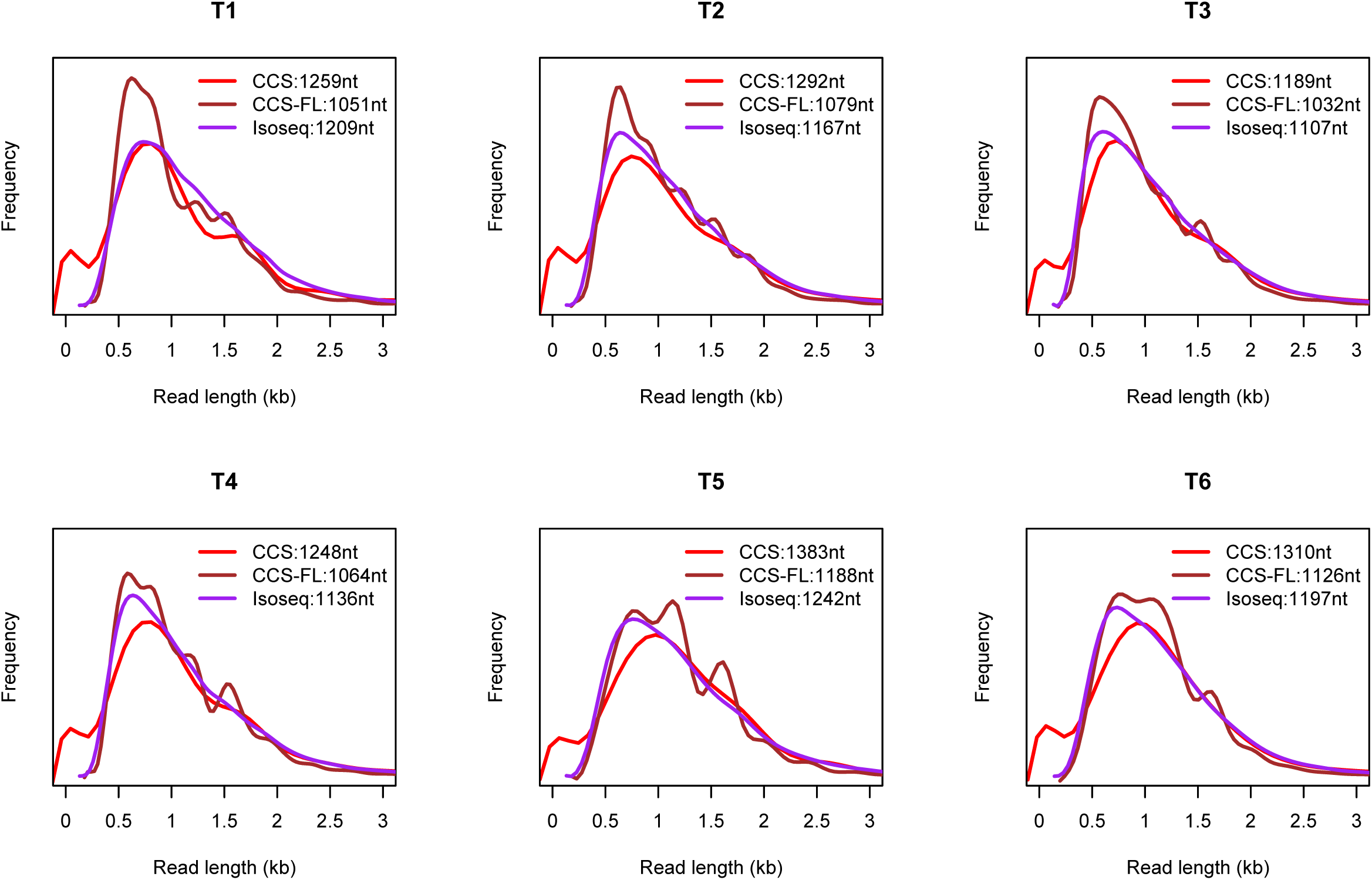
Length distribution of CCS, CCS-FL and Isoseq reads at the 6 different time points.

**Figure S3.**
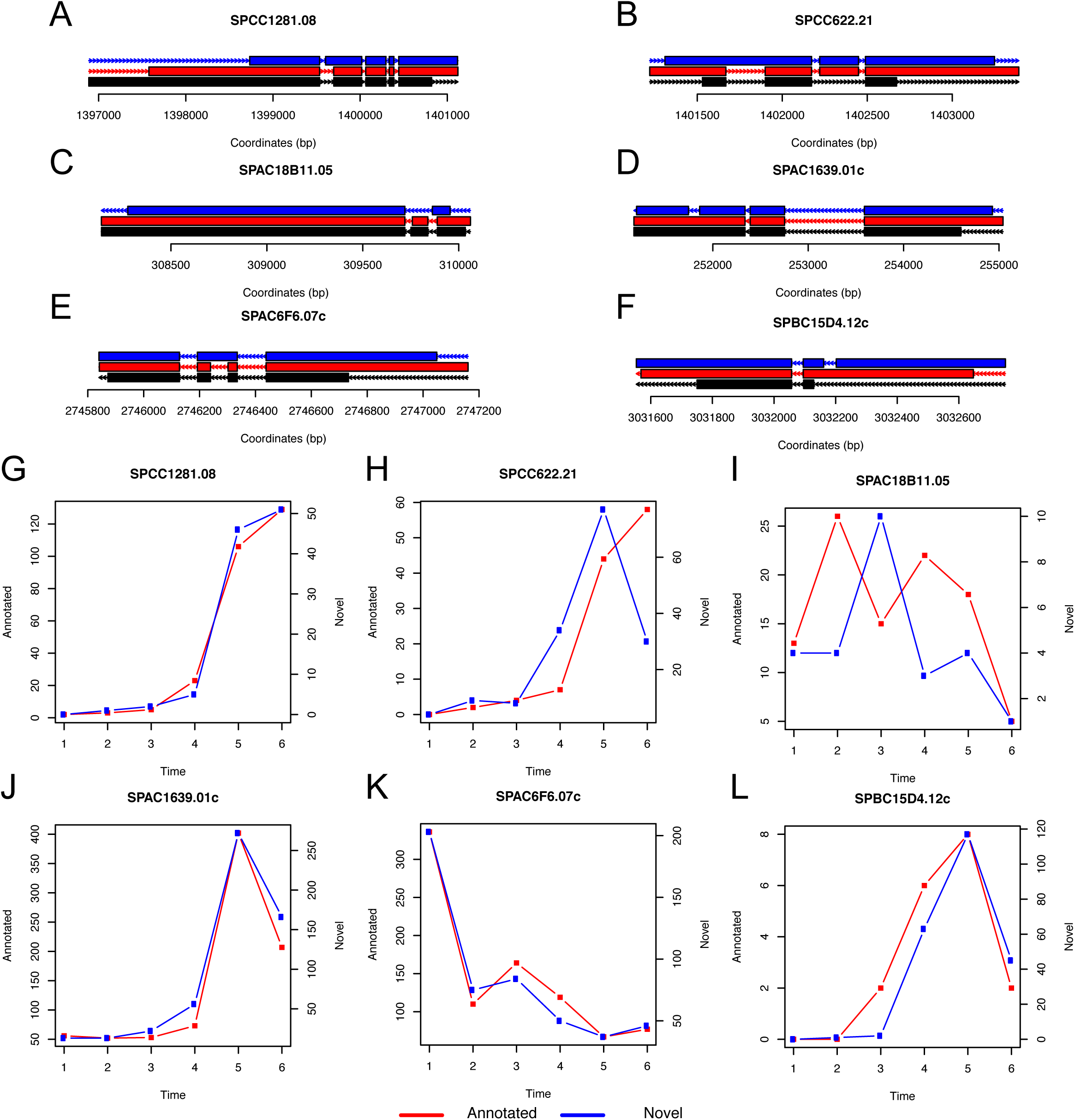
Examples of alternative splicing. (A-F) shows 6 different types of alternative splicing: novel splicing donor or acceptor, exon inclusion, exon skipping, intron in exon, intron retention, novel exon. (G-L) shows the temporal patterns of the annotated and alternative isoforms.

**Figure S4.**
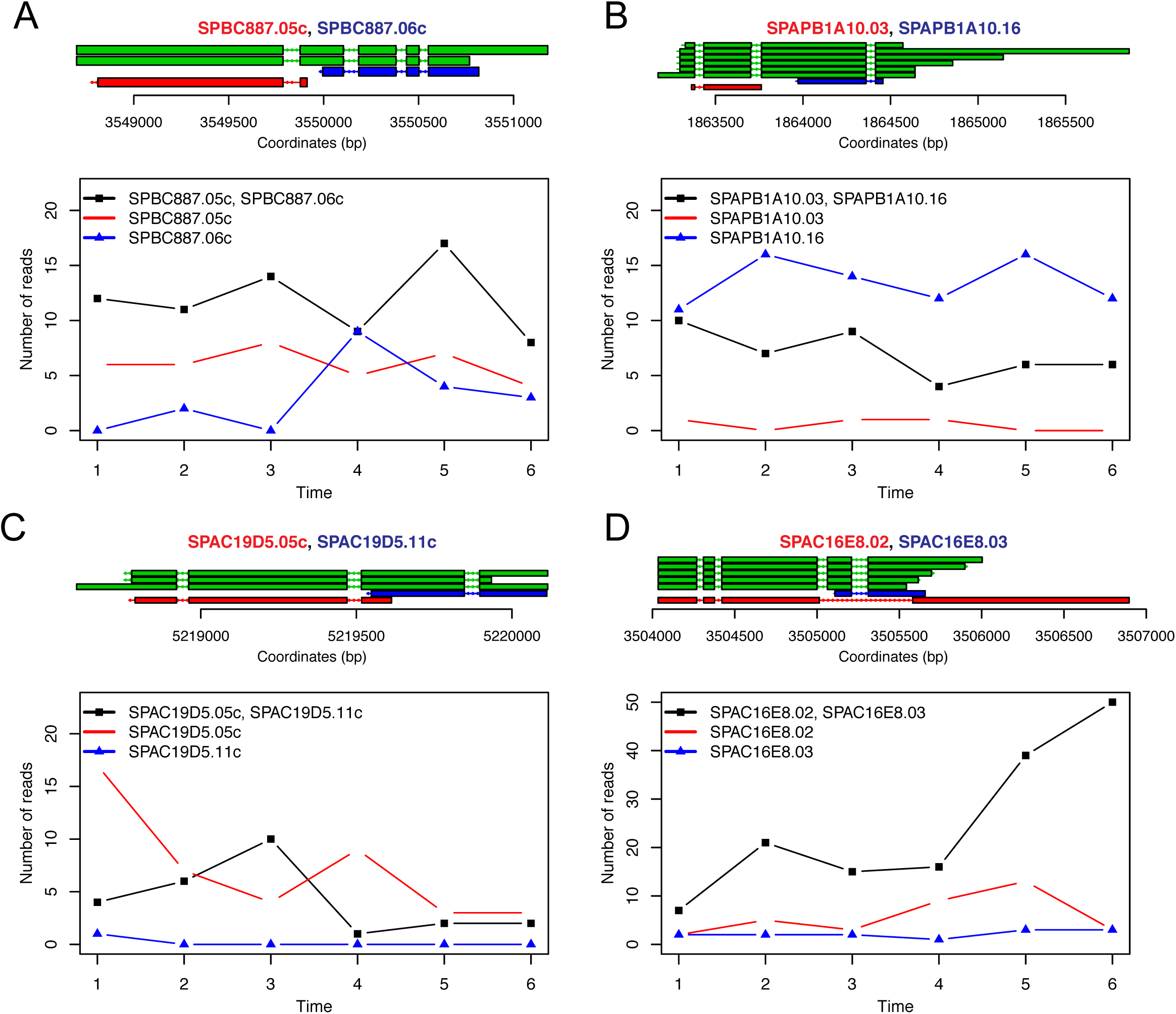
Structures and dynamics of read-through transcripts. (A) shows SPBC887.05c, SPBC887.06c. (B) shows SPAPB1A10.03, SPAPB1A10.16. (C) shows SPAC19D5.05c, SPAC19D5.11c. (D) shows SPAC16E8.02, SPAC16E8.03. The top panels show the structures of isoforms and the bottom panel shows the dynamics of read-through transcripts and corresponding individual transcripts.

**Figure S5.**
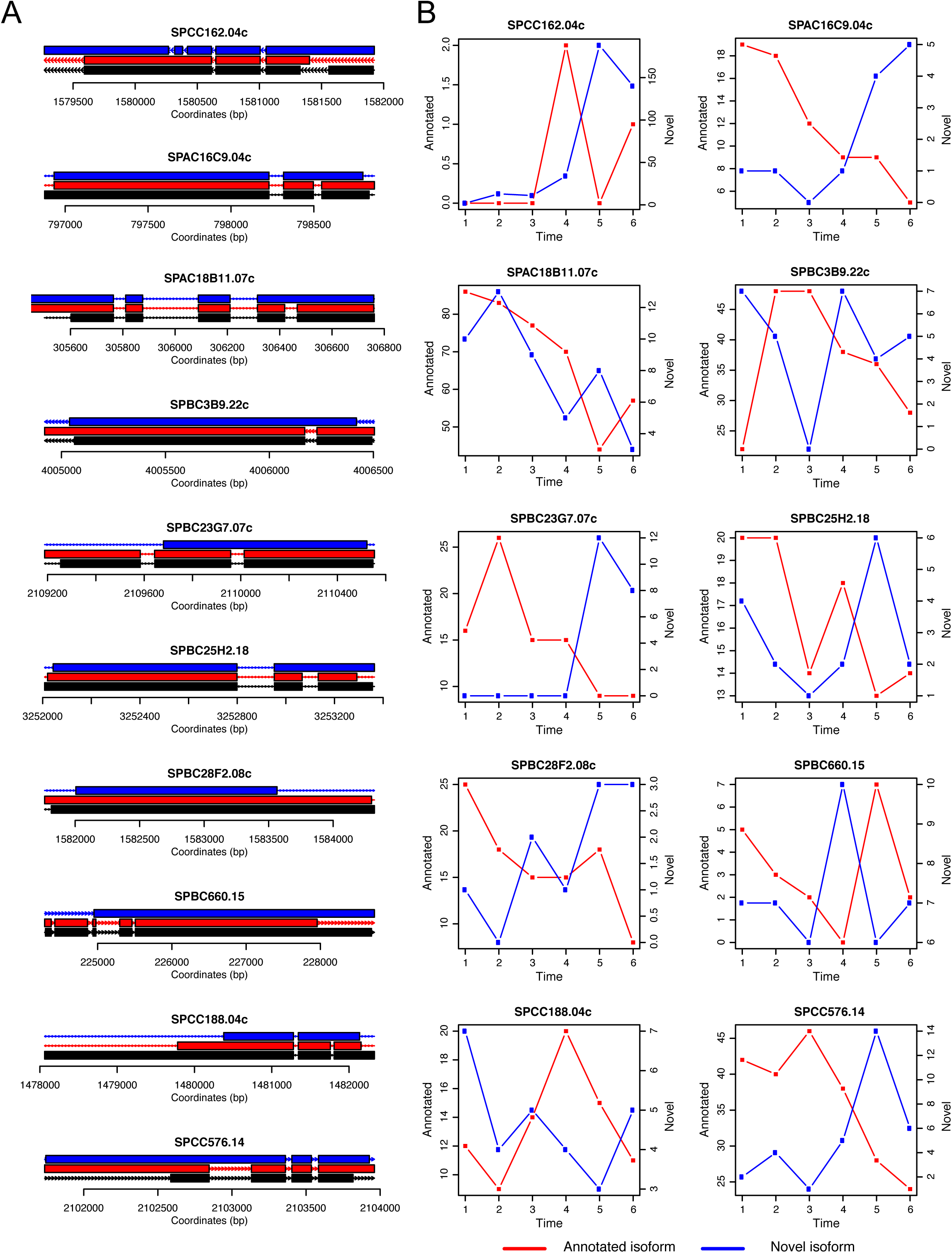
Examples of anticorrelated isoforms in meiosis. (A) shows the exon-intron strucutres of the examples. (B) shows the temporal patterns of the annotated and alternative isoforms.

**Figure S6.**
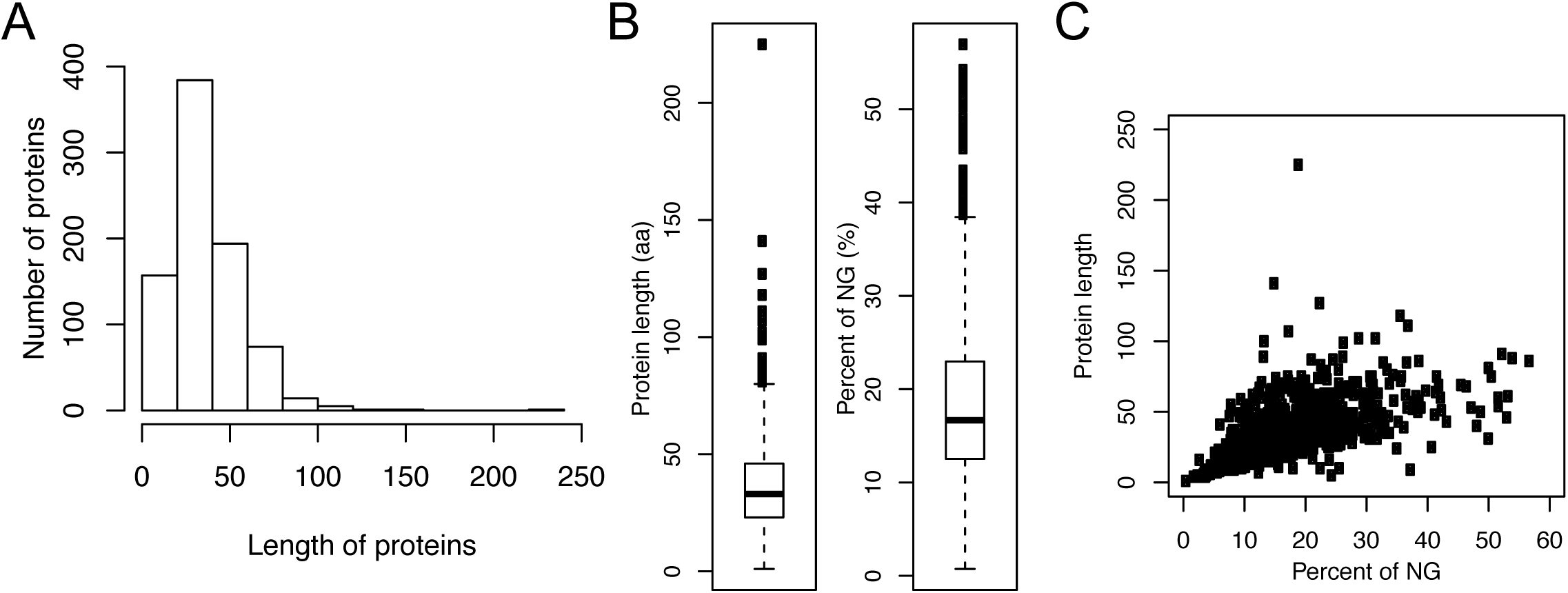
Prediction of translational products from newly discovered transcripts. (A) Distribution of protein length predicted from novel genes. (B) Boxplots showing the length of protein and the percent of novel genes used for translation. (C) Scatterplot of the percent of new genes used for translation and the length of proteins.

**Figure S7.**
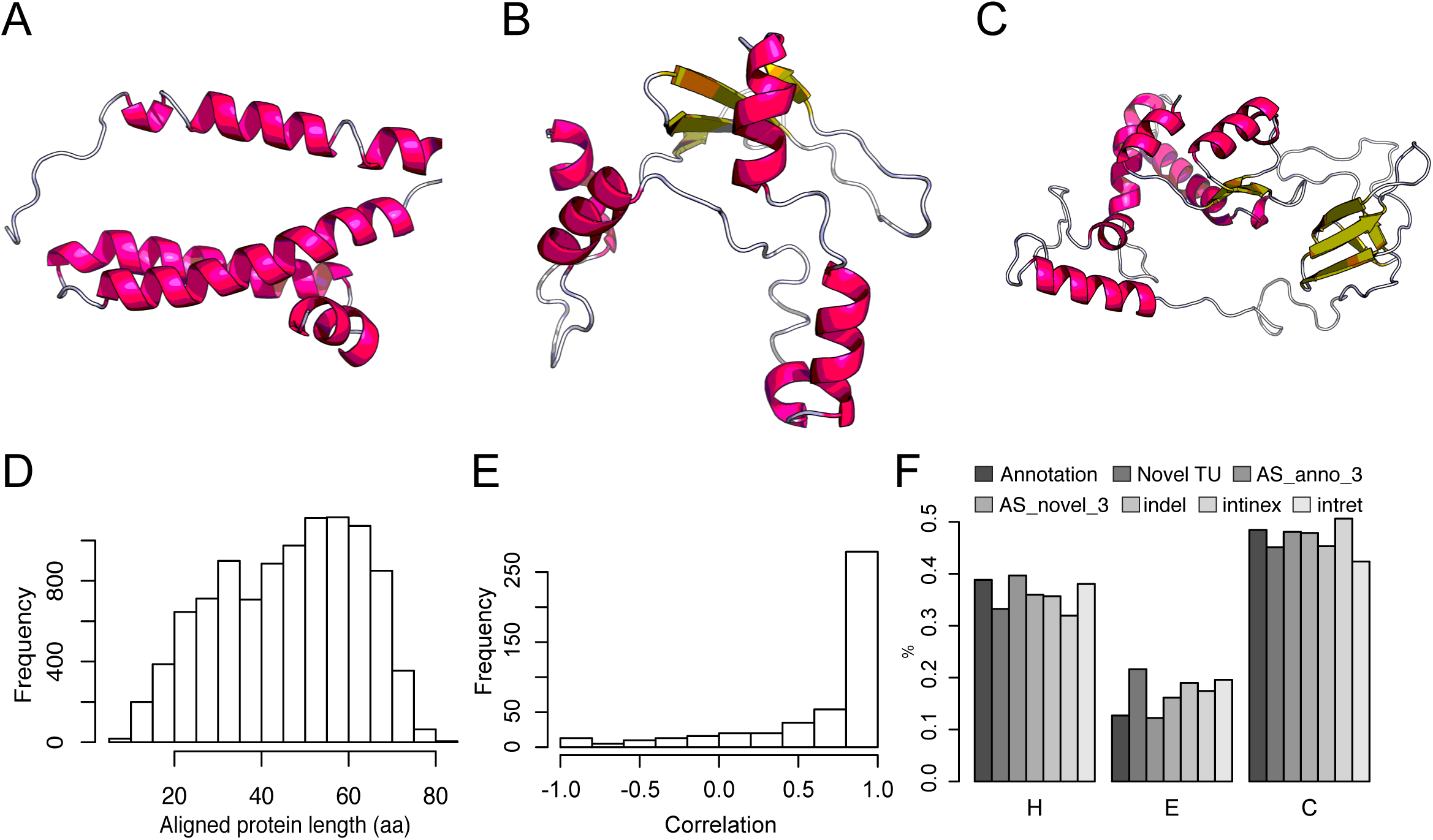
Secondary structure prediction. (A) structure predicted for NG629. (B) structure predicted for NG634. (C) structure predicted for NG653. (D) Lengths of aligned proteins predicted from novel TUs. (E) Correlation of the ratios of α-helix (H), β-sheet (E) and coil (C) predicted from C terminus annotated or amino acid sequence starting at first alteration in AS isoforms. (F) Percentages of α-helix (H), β-sheet (E) and coil (C) predicted for different classes of proteins.

## Table legend

Table S1. Summary of isoforms. The 8^th^ column is the number of CCS FL reads by each replicate at each time point. The 9^th^ column summarizes the type of each AS event along a given isoform as well as their impact on the protein sequence level. From an alignment of altered and annotated protein sequences, the number of inserted or deleted amino acids and the resulting shift in reading frame is depicted. The format representing the alignment is described in detail in Methods. The 10th column gives the multiplicity of each AS type.

Table S2. Summary of numbers of reads.

Table S3. Numbers of AS events for different cutoffs of supporting CCS FL reads.

Table S4. Number of isoforms per gene for different cutoffs of supporting CCS FL reads.

Table S5. Number of CCS FL reads supporting annotated, novel and antisense isoforms.

Table S6. Antisense isoforms.

Table S7. Summary of dinucleotides at novel splicing junctions.

Table S8. Summary of read-through transcripts.

Table S9. Number of CCS FL reads supporting annotated, novel and antisense isoforms at each time points.

Table S10. FASTA file of predicted protein sequences.

Table S11. Translational status. “Both” means this gene may generate both annotated and novel proteins. “WT” means this gene only generates annotated proteins. “Novel” means this gene may only generate novel proteins.

Table S12. Summary of BLAST results for novel TUs and novel AS isoforms.

